# Molecular Mechanisms Associated With Amelioration Of Radiation Induced Gastrointestinal Mucositis By Compound Kushen Extracts

**DOI:** 10.64898/2026.01.12.698969

**Authors:** Yan Zhou, Yuka Harata-Lee, Zhipeng Qu, Hanyuan Shen, Yameng Zhang, Xiumei Duan, David L. Adelson

**Affiliations:** School of Biological Sciences, Faculty of Sciences, Engineering and Technology, Adelaide University, Adelaide, SA, Australia; Zhendong Research Institute, Zhendong Pharmaceutical, Beijing, China; South Australian Museum, North Terrace, Adelaide, SA 5000, Australia

**Keywords:** Gastrointestinal mucositis, Radiotherapy, Compound Kushen Injection, Compound Kushen Powder, Transcriptomic analysis

## Abstract

Radiation induced gastrointestinal mucositis (GIM) is a severe complication of radiotherapy that compromises patient quality of life and treatment efficacy. This study assessed the therapeutic potential and molecular mechanisms of two forms of herbal extracts, Compound Kushen Powder (CKP) and Compound Kushen Injection (CKI) in a rat model of GIM. Administration of either CKP or CKI to irradiated animals significantly reduced the severity of GIM symptoms. Transcriptomic analysis of jejunum and colon mucosa identified candidate mechanisms as well as common pathways between CKP and CKI. CKP modulated innate immune activation and inflammatory signalling, affecting genes such as *Socs3* and *Tlr4*, while CKI promoted tissue repair and oxidative stress resistance through genes including *Osm, Epha2*, and *Stat3*. However, both forms of herbal extract enhanced stress response and stimulus regulation in our GIM model. These findings reveal common mechanisms shared by CKP and CKI that suppress symptoms of radiation induced GIM, along with extract specific effects on distinct pathways, acting on different set of genes.

## 1. Introduction

GIM is a severe complication commonly observed in cancer patients undergoing radiotherapy. It presents a variety of symptoms, including abdominal pain, diarrhoea, and increased intestinal permeability, significantly impacting patient quality of life. Clinically, gastrointestinal symptoms, such as diarrhea and abdominal pain, are frequently observed in 20–70% of patients undergoing radiotherapy for abdominal or pelvic malignancie (Classen et al., 1998). This radiation-induced toxicity is a particular concern in gynecological cancers, affecting approximately 47% of women treated for cervical or endometrial cancer (Abayomi et al., 2009). Moreover, the burden of symptoms extends to other malignancies; up to 80% of patients with head and neck cancer reportedly experience similar complications during radiotherapy (Rubenstein et al., 2004).

The pathogenesis of GIM is complex and involves a cascade of molecular and cellular events. As we have recently described (2024), radiation exposure initially induces DNA damage in epithelial and submucosal cells, leading to an accumulation of reactive oxygen species (ROS) and inflammatory cytokines. These factors trigger downstream signalling pathways, amplifying inflammation and apoptosis of those cells. The consequent epithelial barrier disruption allows bacterial colonisation, further exacerbating inflammation and tissue damage. Under these conditions, intestinal mucositis develops, contributing to pain and gastrointestinal dysfunction (Zhou et al., 2024). Severe cases often lead to treatment interruption, prolonged hospitalization, and increased healthcare burdens, highlighting the urgent need for effective therapeutic interventions.

Despite the significant clinical impact of GIM, its underlying molecular mechanisms remain incompletely understood. The widely accepted five-phases of mucositis progression consist of (1) initiation, (2) upregulation and message generation, (3) signalling and amplification, (4) ulceration and inflammation, and (5) healing (Logan et al., 2007). During initiation, radiation-induced DNA damage results in cellular apoptosis and the production of ROS, activating inflammatory pathways such as those mediated by p53 and nuclear factor kappa-B (NF-κB) (Criswell et al., 2003). The subsequent upregulation and message generation phase further amplifies inflammation through the activation of NF-κB, leading to increased expression of pro-inflammatory cytokines, including tumour necrosis factor (TNF), interleukin-6 (IL-6), and interleukin-1β (IL-1β) (Sonis, 2002). The amplification phase strengthens this inflammatory cascade and creates a positive feedback loop, promoting apoptosis and further tissue damage. During ulceration, the damaged intestinal epithelium provides ground for bacterial colonization, aggravating inflammation and immune activation (Sonis, 2007; Stringer and Logan, 2015). The final healing phase involves epithelial migration and tissue repair, but the regenerative process may be delayed due to persistent inflammatory stress (Tran et al., 2004).

As GIM continues to be a significant unmet medical need due to the lack of clinically effective therapies, improving patient outcomes and reducing treatment-related morbidity require innovative therapeutic approaches that target its biological processes. Compound Kushen Injection (CKI) is a traditional Chinese medicine formulation derived from *Radix Sophorae Flavescentis* (Kushen) and *Rhizoma Smilacis Glabrae* (Baituling). It is primarily indicated for the treatment of cancer-related pain and bleeding (Shen et al., 2019). Its efficacy as an anti-inflammatory and analgesic provide continued research interest into its potential therapeutic applications. CKI contains active compounds such as matrine and oxymatrine, which have been shown to modulate TRPV1-mediated signalling and suppress pro-inflammatory cytokine secretion (Zhao et al., 2014). Additionally, CKI enhances antioxidative defences by regulating glutathione metabolism and modulating immune responses through the PI3K/Akt pathway (Wang et al., 2023). It also modulates ROS levels, indicating an antioxidative capacity by reducing intracellular ROS (Jin et al., 2018). In our laboratory, a rat model of radiation-induced GIM was established to assess efficacy of CKI in reducing severity of mucositis and to elucidate its underlying mechanisms. The study showed that CKI administration resulted in significantly less damage to intestinal villus epithelium, a reduction in apoptotic cells within intestinal crypts, and decreased levels of inflammatory markers such as interleukins IL-1β, IL-6, and myeloperoxidase-producing cells (Harata-Lee et al., 2022).

Building on these findings, Compound Kushen Powder (CKP) has been developed as an oral formulation aimed at improving treatment accessibility and efficacy. While CKI has been well studied in injectable forms, local bioavailability of CKP and its potential to mitigate radiation-induced mucosal damage remain unexplored. Given its phytochemical composition, CKP is believed to have similar anti-inflammatory and antioxidant effects as CKI, making it a promising candidate for alleviating GIM symptoms.

## 2. Materials and Methods

### 2.1 Compound Kushen preparation and administration

Both Compound Kushen Preparation (CKP) and Compound Kushen Injection (CKI, Batch No. 20210519) were provided by Zhendong Pharmaceutical Co. Ltd. (Shanxi, China). CKP (containing 71.81 mg/g total alkaloids) was suspended in sterile water at 0.28 g/ml to achieve a final concentration of 20 mg/ml. The vehicle control for CKI was prepared using 0.25% Tween 80 (#P6249, Sigma Aldrich, MO, USA) and 25 mM HEPES (#15630080, Thermo Fisher Scientific, MA, USA) in sterile H_2_O. CKP was delivered via oral gavage, whereas CKI was administered via intraperitoneal injection. Both treatments (or their respective vehicles) were administered at doses of 100 mg/kg (5 ml/kg) or 140 mg/kg (7 ml/kg) of total alkaloids. All treatments were given daily from day 0 to day 6, 2 hours prior to irradiation.

### 2.2 Animal model and irradiation protocol

All experimental procedures were approved by the Animal Ethics Committee at South Australia Health and Medical Research Institute (SA, Australia). Eight-week-old male Sprague-Dawley rats were previously used to establish a GIM model (Harata-Lee et al, 2022). Briefly, rats were anaesthetised, placed in lead shielding capsule, and receive fractionated abdominal irradiation at 4 Gy/day for 5 consecutive days (total dose 20 Gy). The study included four experimental groups: (1) water vehicle, (2) CKP, (3) Tween 80/HEPES vehicle, and (4) CKI. GIM progression was assessed daily through monitoring of weight, food intake, and diarrhoea severity. Diarrhoea severity was scored using a scale from 0 to 3 in increments of 0.5 as previously described (Harata-Lee et al., 2022).

### 2.3 Sample collection and processing

On days 4 and 7 post irradiation, small and large intestines were removed and rinsed with cold saline solution. Approximately 1 cm segments were excised from the jejunum and colon, then immediately frozen in liquid nitrogen for storage. Mucosal layers were collected from the tissues for RNA extraction with the PureLink RNA Mini Kit (Thermo Fisher Scientific). RNA quantity and quality were first checked using a NanoDrop 2000 spectrophotometer (Thermo Fisher Scientific). To ensure RNA integrity, further testing was done with the Agilent 2100 Bioanalyzer (Agilent Technology, CA, USA) at SA pathology (SA, Australia). RNA samples with RIN>6.0 were submitted to BGI Genomics (Hong Kong, China) for strand-specific mRNA library construction after poly(A) selection and 150 bp paired-end sequencing on the DNBSEQ platform.

### 2.4 Transcriptome data analysis and functional annotation

Raw sequencing reads were quality-controlled using FastQC (v0.11.9). Adapters and low-quality bases were trimmed with bbduk (BBMap version 39.01) under the parameters: hdist=1 mink=11 ktrim=r qtrim=rl trimq=20 minlen=25 trimpolyg=20. Clean reads were aligned to the *Rattus norvegicus* reference genome (mRatBN7.2) using STAR (v2.7.10a) with the parameters: --outFilterMismatchNoverLmax 0.03 and --alignIntronMax 10000. Low-expression genes which didn’t pass a threshold of Counts Per Million (CPM) > 1 in at least 50% of samples were filtered. Normalization and quality control were performed using the “voom” method from the “limma” package, ensuring sample-specific adjustments (Liu et al., 2015). Differentially expressed (DE) genes were identified with thresholds of |log2 fold change| > 1 and false discovery rate (FDR) < 0.05 using limma-trend approach of the “limma” package. Functional enrichment analysis was performed using the DAVID Gene Ontology (GO) and Kyoto Encyclopedia of Genes and Genomes (KEGG) pathway databases. Enrichment analysis thresholds were set at a minimum gene count of 5 and a maximum EASE score of 0.05.

### 2.5 Statistical analysis

Statistical analysis was conducted using GraphPad Prism version 9.0.2 (GraphPad Software, San Diego, CA, USA). Data were presented as mean ± standard deviation of the mean (SD). For datasets plotted over time (e.g., diarrhoea severity, represented as XY line and bar graphs), a two-way analysis of variance (ANOVA) was applied to compare the treatment effects of CKP and CKI against the vehicle control group. A *P*-value < 0.05 was considered statistically significant.

## 3. Results

### 3.1 CKP and CKI mitigate radiation-induced diarrhoea in GIM

Consistent with our previous findings that concurrent treatment effectively reduces GIM severity whereas post-irradiation treatment does not, a concurrent treatment dosing plan was applied in this study (Harata-Lee et al., 2022). Time-course analysis of diarrhoea severity in irradiated rats demonstrated a marked increase in diarrhoea scores following abdominal irradiation, peaking on day 7 post-irradiation (Figure 1A). Compared with the vehicle control group, both 100 mg/kg and 140 mg/kg CKP treatment groups showed a significant reduction in diarrhoea scores (Figure 1A). To further investigate treatment effects at different disease phases, diarrhoea scores were averaged for the onset phase (days 0–7) and recovery phase (days 7–11). CKP groups significantly relieved diarrhoea symptoms during day 0 to day 7, compared to the vehicle control (Figure 1B). A focused comparison of diarrhoea scores at peak severity (day 7) confirmed this trend with both 100 mg/kg CKP and 140 mg/kg CKP groups showing a significant reduction in diarrhoea scores compared to the vehicle control (Figure 1C). Because diarrhoea can cause dehydration and nausea/loss of appetite, we measured body weight loss as well. As shown in Figure 1D, CKP administration effectively mitigated radiation-induced weight loss.

**Figure 1:**
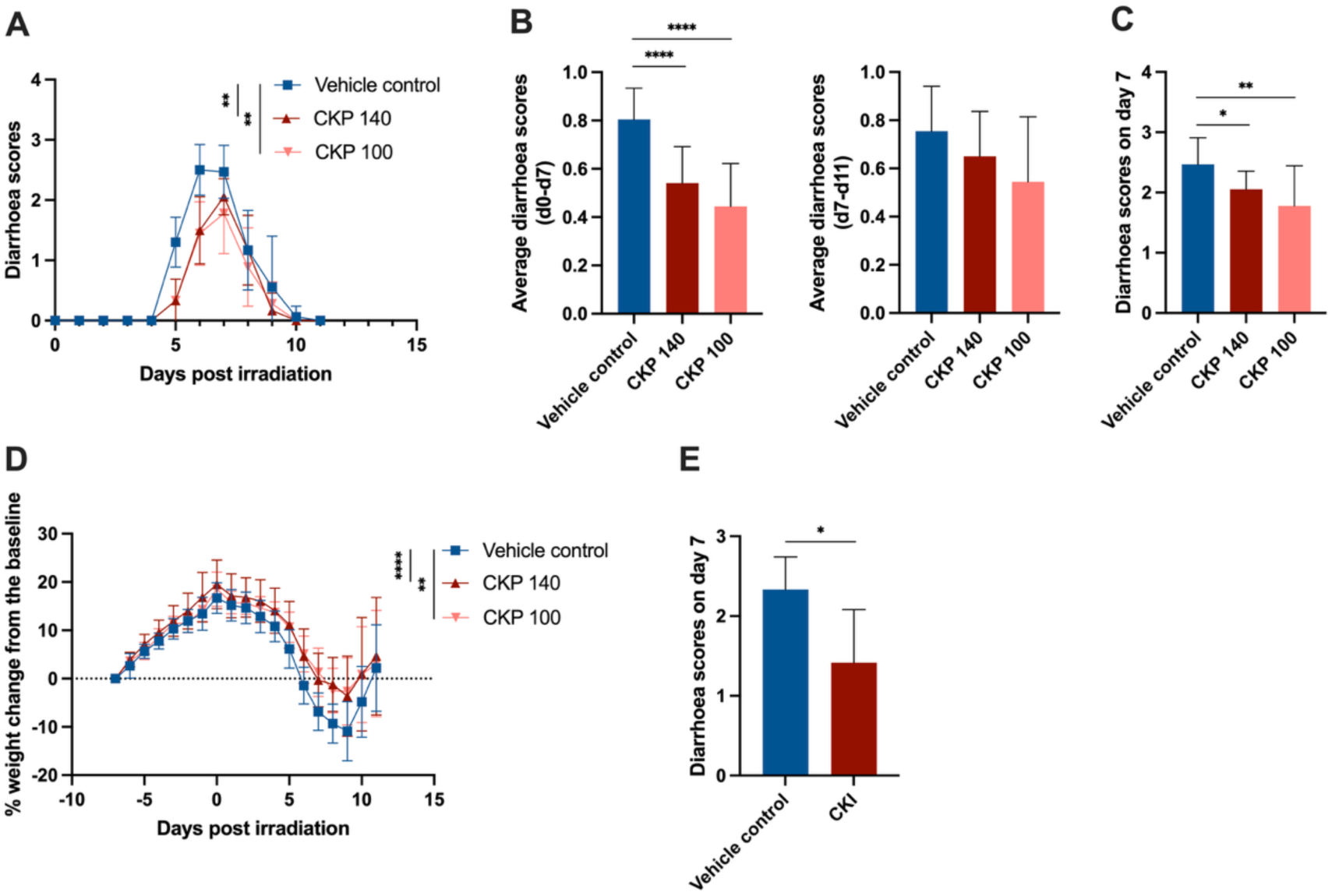
Effect of CKP or CKI on severity of radiation induced GIM. (**A**) Diarrhea scores of irradiated Sprague-Dawley rats. Rats received five daily fractional doses of radiation at 4 Gy/day (20 Gy total) on their abdomen and were concurrently treated with CKP or vehicle control daily. The animals were monitored daily for the presence and degree of diarrhea and given a score according to the scoring scheme (see Methods). (**B**) Comparison of average diarrhoea scores during the onset (left) and the recovery (right) phases. (**C**) Comparison of diarrhoea scores at the peak of the symptoms. (**D**) Body weight changes of irradiated rats treated with CKP or vehicle control. (**E**) Diarrhoea scores of irradiated rats that were treated with CKI or vehicle control. Data are presented as mean±SD.

Our previous study indicated that CKI doses of 2 ml/kg and 3 ml/kg can significantly mitigate diarrhoea symptoms (Harata-Lee et al., 2022). A direct comparison of CKI with vehicle control was carried out using a dosage of 2.5 mL/kg CKI to evaluate its therapeutic effects, with diarrhoea scores assessed on day 7 following irradiation. CKI treatment significantly reduced diarrhoea scores compared to the vehicle control (Figure 1E).

### 3.2 Molecular responses to CKP treatment in radiation-induced GIM

To investigate the molecular mechanisms of CKP and CKI in mitigating radiation-induced GIM injury, transcriptomic data from jejunum and colon epithelial tissues collected at the onset and peak of the disease course were analysed. The eight comparisons are summarised in the table, detailing treatment groups, tissue types, timepoints and corresponding vehicle controls (Figure 2). The initial comparison of gene expression data for each drug at different time points in two different tissues were carried out with respect to their vehicle controls. The comparisons of DE genes from CKP treated groups against vehicle controls revealed distinct variations across different tissues and time points. Specifically, colon tissue on day 4 had 2035 DE genes, decreasing on day 7 to 174 DE genes. Jejunum tissue on day 4 had 453 DE genes decreasing on day 7 to 320 DE genes (Supp Figure 1). Comparison of DE genes from the above four DE gene sets showed that only one gene was shared across all four groups (Figure 3A). When comparing the DE genes identified from the two different tissues at the same time point, 62 common genes were observed in both the Colon and Jejunum groups at day 4, and 6 genes were observed in both tissues at day 7 (Figure 3A). When comparing the DE genes identified from the two different time points in the same tissue, 29 common genes were colon-specific at both day 4 and day 7, and 15 common genes were observed as jejunum-specific at both time points (Figure 3A).

**Figure 2:**
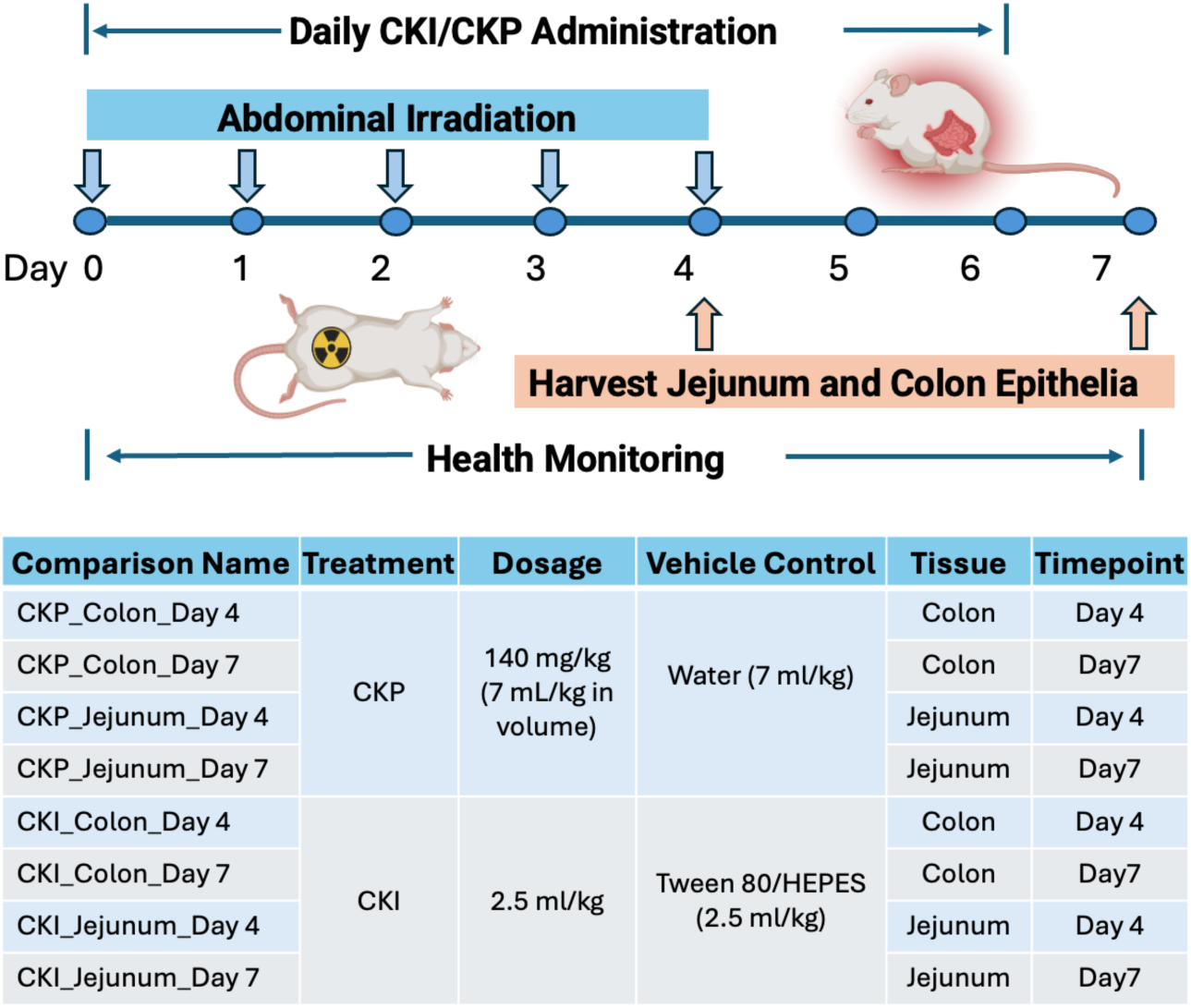
Experimental design for the transcriptome analyses. Rats received abdominal irradiation from day 0 to day 4. Daily administration of CKI, CKP, or vehicle control continued from day 0 to day 6. Each group included six rats and health monitoring was conducted throughout the study. Rats were euthanized on day 4 or day 7 post-irradiation, and jejunum and colon epithelial tissues were collected for transcriptomic analysis.

**Figure 3:**
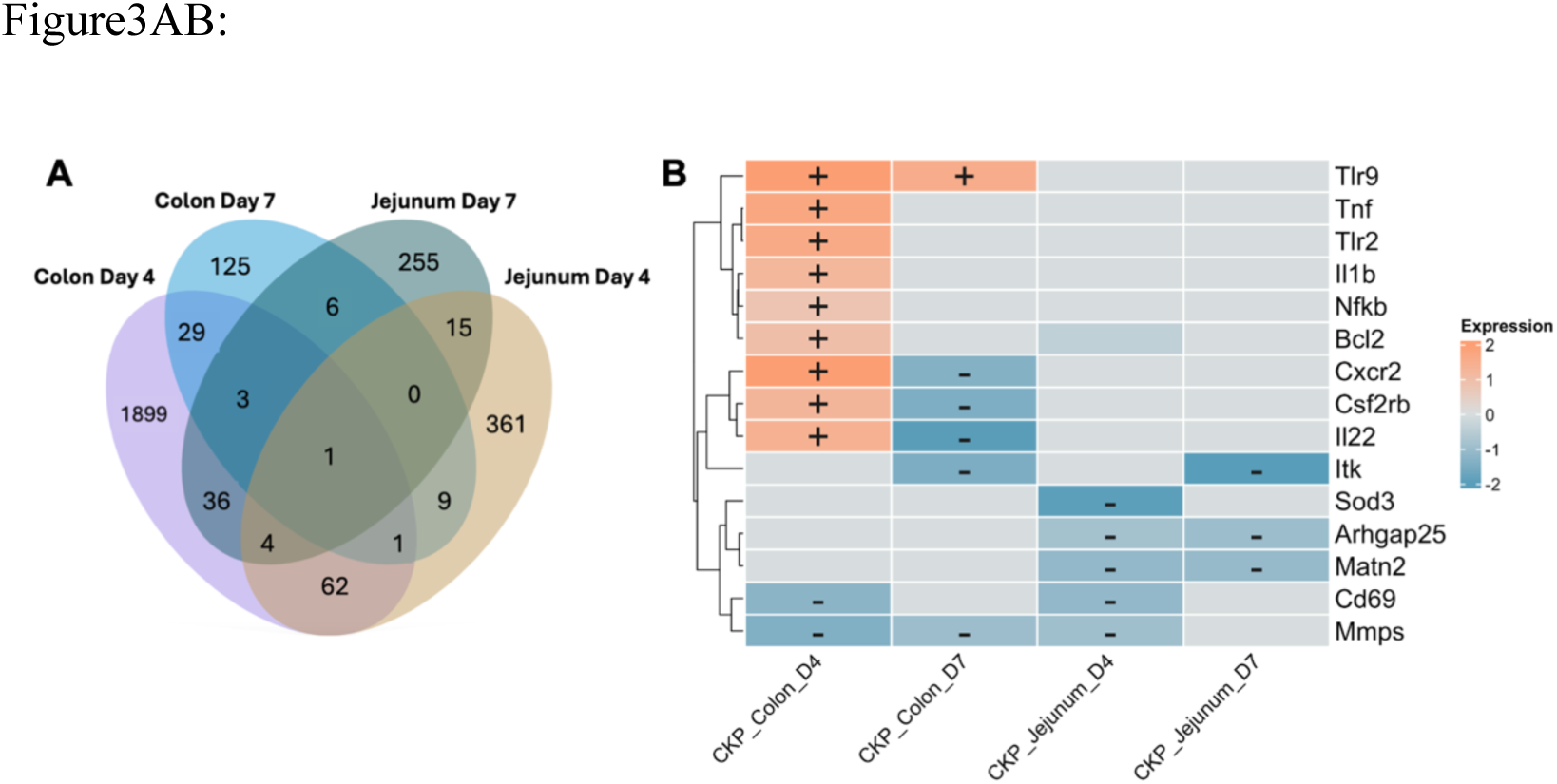
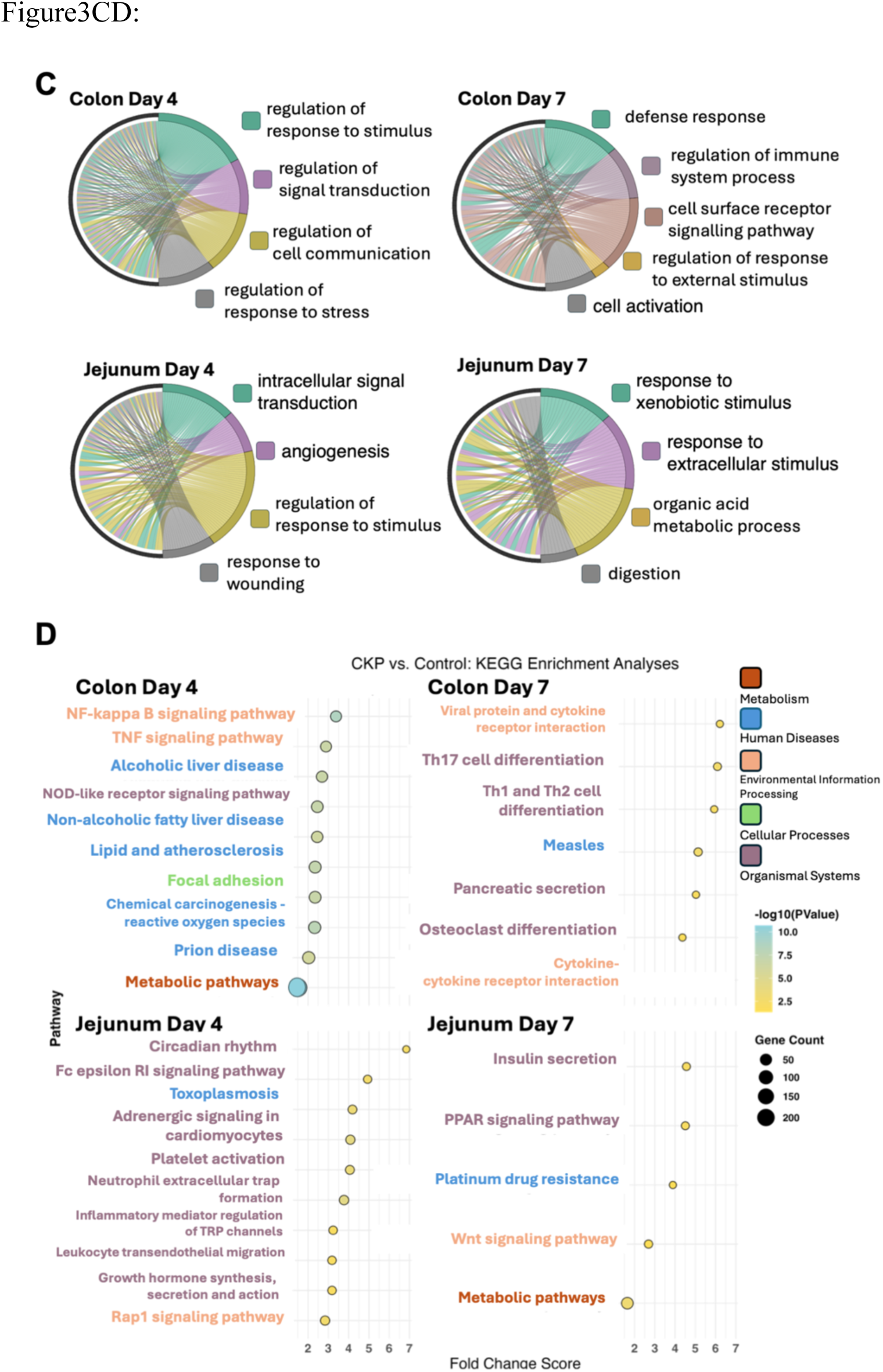
Transcriptomic analysis of CKP-treated tissues. (**A**) Venn diagram showing the numbers of DE genes between CKP and vehicle control treated colon and jejunum at day 4 and day 7 of irradiation. Each region represents unique and shared DE genes between the different comparisons. (**B**) Heatmap of mucositis-related DE genes in CKP-treated colon and jejunum at day 4 and day 7 post-irradiation. Orange grids indicate positive fold change, while blue represents negative fold change. (**C**) GO enrichment analysis of DE genes in CKP-treated tissues visualized using chord plots. Different colours represent distinct biological processes, with line density indicating the number of genes associated with each GO term. Genes are arranged anti-clockwise by logFC from highest to lowest on the left side of the plot; therefore, GO terms connected to DE genes clustered near the 12-o’clock position correspond to biological processes involving genes with the largest expression changes in this comparison. (**D**) KEGG pathway enrichment analysis of DE genes in CKP-treated colon and jejunum at day 4 and day 7. The plots display the top enriched pathways ranked by Fold Changed Scores, with dot size representing the number of genes involved and colour intensity indicating statistical significance. Pathways are classified according to KEGG Level 1 categories, represented by distinct colours.

To further investigate the molecular effects of CKP on GIM, we focused on a curated set of 15 mucositis-related genes involved in inflammatory signalling, immune modulation, and tissue repair, which are key processes in the pathogenesis and recovery of GIM (Keefe et al., 2004; Korpos et al., 2015; Radulovic and Niess, 2015; Shen et al., 2018, 2018; Stadnicki, 2011; Zhou et al., 2024). The selected genes include Toll-like receptor 9 (*Tlr9*), Interleukin-22 (*Il22*), colony-stimulating factor 2 receptor subunit β (*Csf2rb*), C-X-C motif chemokine receptor 2 (*Cxcr2*), Rho GTPase activating protein 25 (*Arhgap25*), Matrilin-2 (*Matn2*), CD69 molecule (*Cd69*), IL-2 inducible T-cell kinase (*Itk*), B-cell lymphoma 2 (*Bcl2*), matrix metalloproteinases (*Mmps*), Tumour necrosis factor α (*Tnf*), Interleukin-1 beta (*Il1b*), Nuclear factor κB (*Nf*-*kb*), Toll-like receptor 2 (*Tlr2*) and Superoxide dismutase 3 (*Sod3)*. A heatmap was generated to visualize the expression patterns of these genes in colon and jejunum tissues for the comparisons between CKP and vehicle control treatment at day 4 and day 7 post-irradiation (Figure 3B). *Tlr9, Tnf, Tlr2, Il1b, Nf-kb, Bcl2, Cxcr2, Csf2rb*, and *Il22* were significantly upregulated under the treatment of CKP in colon tissue at day 4, but only one of them, *Tlr9,* was still shown as up-regulated in colon tissue at day 7. Compared to day 4, the majority of the 15 genes in the colon day 7 group showed no significant change or were downregulated. In contrast to the colon, the 15 genes were mostly inhibited (downregulated) in the jejunum.

DE genes enriched in biological process GO terms were analysed and organised in cord plots showing the contribution of genes to the significantly enriched biological processes. In colon tissue on day 4 were primarily related to cellular responses, including regulation of response to stimulus, signal transduction, and cell communication (Figure 3C). At day 7, colon tissue DE genes enriched in biological process GO terms were involved in regulating immune system processes, defence response, and external stimulus regulation. In the jejunum tissue, GO terms with DE genes enriched on day 4 were associated with intracellular signalling transduction, angiogenesis, and wound response. On day 7, the jejunum DE genes enriched in GO terms were linked to metabolic processes, xenobiotic response, and extracellular stimulus regulation (Figure 3C).

KEGG pathway enrichment analysis of DE genes in colon and jejunum tissues from CKP treated groups at days 4 and 7 post irradiation identified distinct biological pathways, and plots were generated to show categories of regulated pathways and magnitude of regulation (Figure 3D). Of those pathways relevant to radiation induced GIM, at day 4 in the colon tissue, multiple pathways were associated with inflammatory signalling, including NF-κB signalling, TNF signalling, and NOD-like receptor signalling. By day 7 in the colon tissue, enriched pathways included Th17 cell differentiation and Th1, Th2 cell differentiation (Figure 3D), which are critical to adaptive immune responses, along with cytokine-cytokine receptor interaction, which contributes to both innate and adaptive immune regulation.

In jejunum tissue at day 4, enrichment analysis revealed several significant pathways including the Rap1 signalling pathway and inflammatory mediator regulation of TRP Channels. By day 7, jejunum pathways were notable by metabolic and repair mechanisms, such as PPAR signalling and *Wnt* signalling. Pathways related to insulin secretion, platinum drug resistance and xenobiotic metabolism were also observed (Figure 3D).

### 3.3 Molecular responses to CKI treatment in radiation-induced GIM

Similarly, comparisons were made between the composition of DE genes and their associated biological functions in the CKI group. Comparison of DE genes in CKI-treated groups against vehicle controls showed distinct transcriptional changes across tissues and time points (Figure 4A). Colon tissue on day 4 exhibited 435 DE genes, compared to 158 DE genes on day 7. In jejunum tissue, 338 DE genes were found on day 4, compared to 218 DE genes on day 7 (Supp Table 2). No genes were shared among all four groups. While there was an overlap of 5-10 genes between each pair of groups. Most of DE genes were unique to each group.

**Figure 4:**
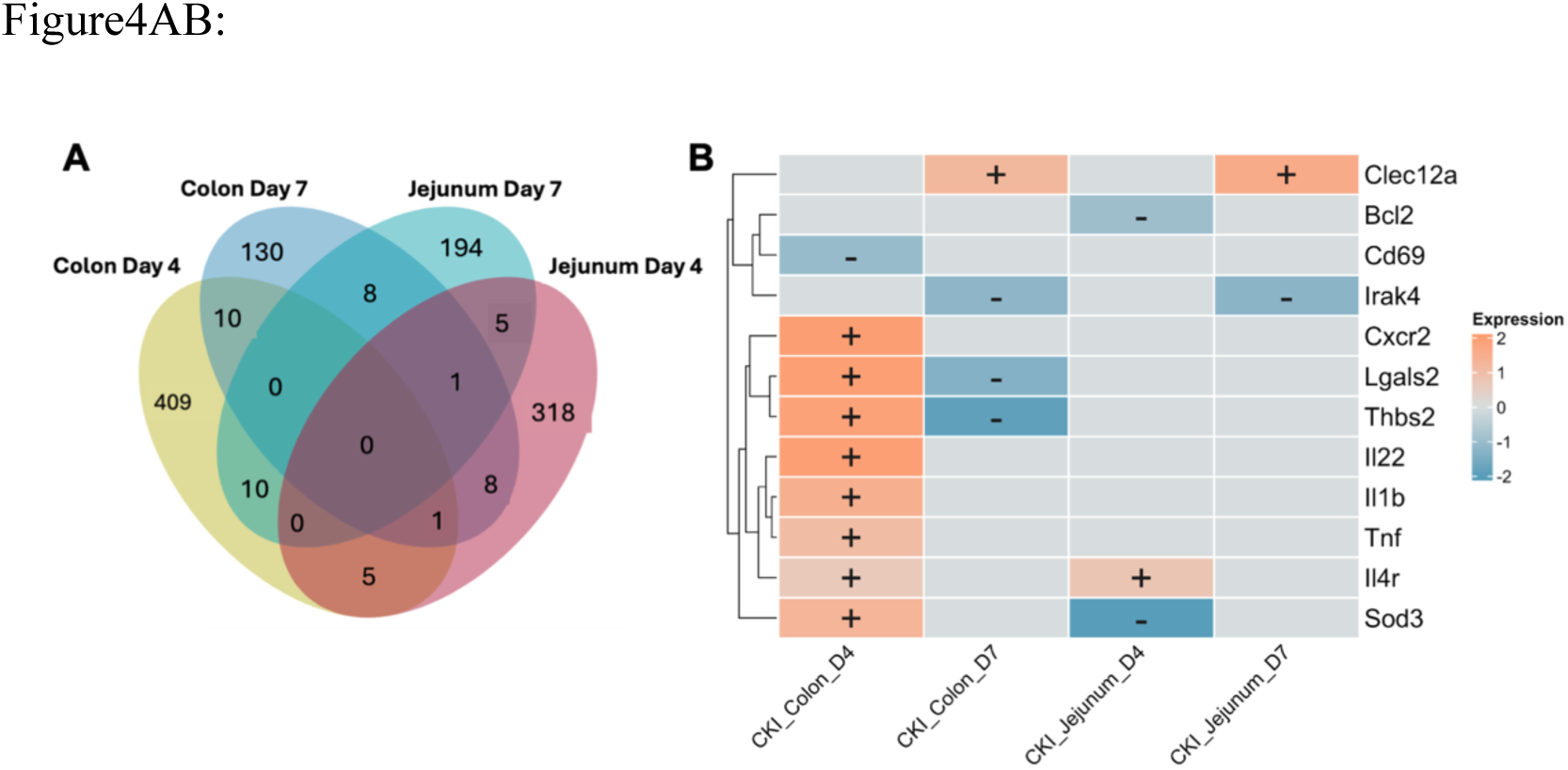
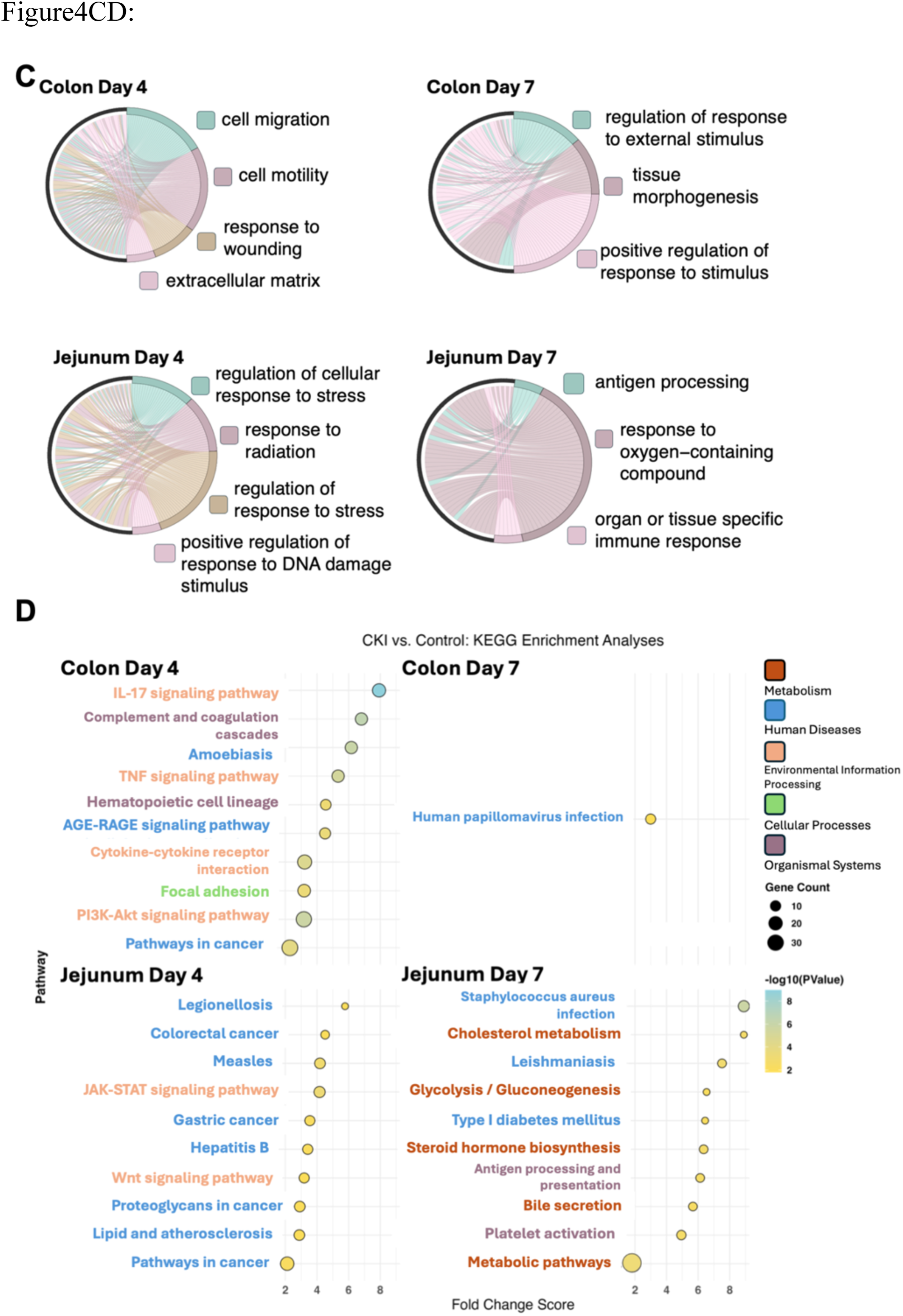
Transcriptomic analysis of CKI-treated tissues. (**A**) Venn diagram showing the numbers of DE genes between CKP and vehicle control treated colon and jejunum at day 4 and day 7 of irradiation. Each region represents unique and shared DE genes between the different comparisons. (**B**) Heatmap of mucositis-related DE genes in CKI-treated colon and jejunum at day 4 and day 7 post-irradiation. Orange grids indicate positive fold change, while blue represents negative fold change. (**C**) GO enrichment analysis of DE genes in CKI-treated tissues visualized using chord plots. Different colours represent distinct biological processes, with line density indicating the number of genes associated with each GO term. Genes are arranged anti-clockwise by logFC from highest to lowest on the left side of the plot; therefore, GO terms connected to genes clustered near the 12-o’clock position correspond to biological processes involving genes with the largest expression changes in this comparison. (**D**) KEGG pathway enrichment analysis of DE genes in CKI-treated colon and jejunum at day 4 and day 7. The plots display the top enriched pathways ranked by Fold Changed Scores, with dot size representing the number of genes involved and colour intensity indicating statistical significance. Pathways are classified according to KEGG Level 1 categories, represented by distinct colours.

To assess the regulatory effects of CKI on GIM, a panel of genes from the identified DE genes was curated based on their relevancy to mucosal injury and repair: Galectin-2 (*Lgals2*), Thrombospondin-2 (*Thbs2*), Sod3, Interleukin-4 receptor α chain (*Il4r*), Interleukin-1 receptor-associated kinase 4 (*Irak4*), C-type lectin domain family 12 member A (*Clec12a*), *Il22*, *Cxcr2*, *Cd69*, *Tnf*, *Il1b*, and *Bcl2* (Espinal et al., 2022; Huang et al., 2024; Keefe et al., 2004; Neumann et al., 2014; Radulovic and Niess, 2015; Shen et al., 2018; Zhou et al., 2024). A heatmap was generated to visualize their expression changes in each set of comparison (Figure 4B). In colon tissues at day 4 post-irradiation, most genes in the panel were upregulated, except for *Cd69*, which was downregulated. By day 7, *Clec12a* remained upregulated, but fewer genes were differentially expressed and only three were down regulated. In jejunum, most genes in the panel didn’t show a change in expression. Only two genes, *Bcl2* and *Sod3*, were down regulated and one gene *Il4r* was up regulated at day 4. At day 7, one gene *Clec12a* was up regulated and one gene *Irak4* was down regulated (Figure 4B).

To explore the biological processes influenced by CKI, GO enrichment analysis of DE genes was also conducted, and chord plots were generated to visualize the findings (Figure 4C). In colon tissue on day 4, the enriched GO terms included cell migration, motility, response to wounding, and extracellular matrix organization. By day 7, these had shifted to response to external stimulus, tissue morphogenesis, and response to external stimuli. In jejunum tissue, day 4 enriched terms were linked to stress responses, including regulation of cellular response to stress, radiation response, and DNA damage repair. By day 7, the enriched terms changed to antigen processing, response to oxygen-containing compounds, and tissue specific immune responses (Figure 4C).

To further understand the molecular mechanisms contributing to CKI mitigation of radiation-induced GIM symptoms, KEGG pathway enrichment of DE genes for comparison between CKI- and vehicle control-treated colon and jejunum tissues at days 4 and 7 post-treatment were examined (Figure 4D). In colon tissue at day 4, pathways associated with inflammatory and immune responses were enriched, including IL-17 signalling, complement and coagulation cascades, TNF signalling, and cytokine-cytokine receptor interaction. Additionally, pathways related to cell adhesion and cancer progression, such as PI3K-Akt signalling and focal adhesion, were identified. By day 7, only human papillomavirus infection pathway was observed, which is not directly linked to GIM modulation. In jejunum, pathways enriched on day 4 were associated with JAK-STAT signalling, *Wnt* signalling, and various cancer-related pathways. By day 7, metabolic and immune-associated pathways became more prominent, including glycolysis/gluconeogenesis, cholesterol metabolism, antigen processing and presentation, and platelet activation (Figure 4D).

### 3.4 Comparative Analysis of CKP and CKI: Shared and Distinct Gene Expression

To better understand the gene expression differences induced by CKI and CKP treatments, we conducted a comparative analysis of DE genes in each comparison group. The aim was to understand unique and shared transcriptional changes associated with each treatment condition. Venn diagrams were generated illustrating the number of unique and overlapping DE genes in jejunum and colon tissues at Days 4 and 7 post-irradiations (Figure 5). Notably, the proportion of shared DE genes across conditions was very low compared to the proportion of unique genes.

**Figure 5.**
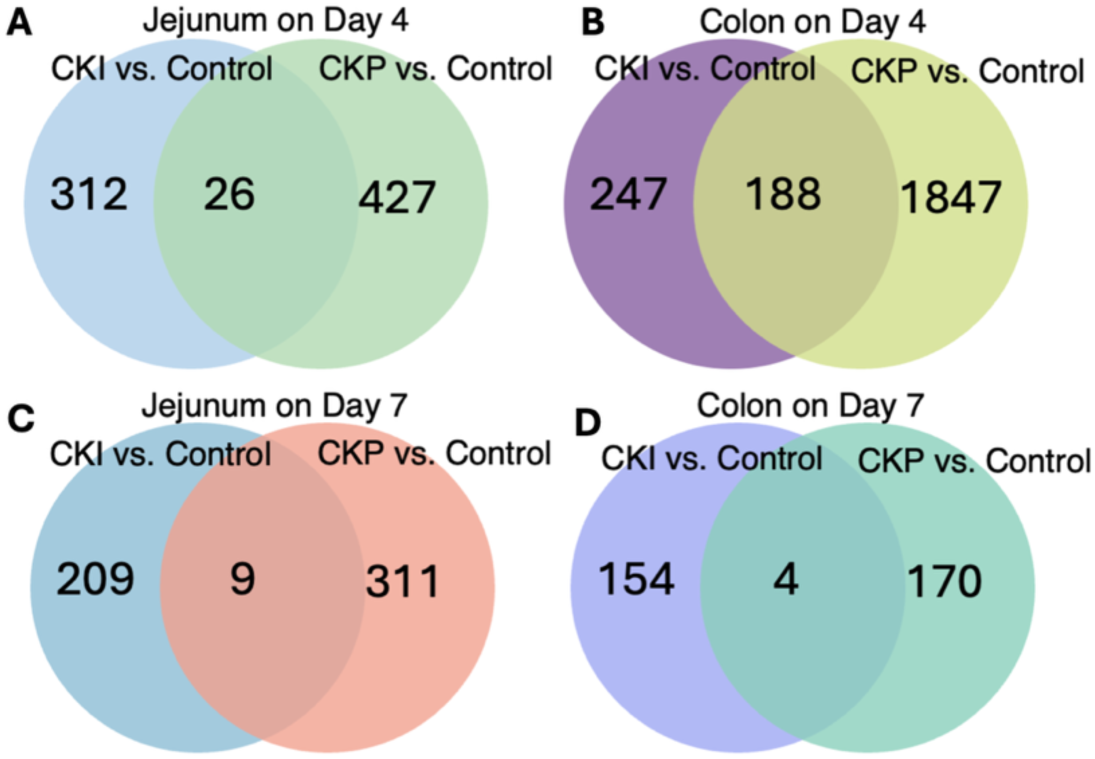
Venn diagrams of DE genes between CKI-treated and CKP-treated tissues. (**A–D**) Venn diagrams showing the unique and overlapping DE genes in jejunum and colon tissues for CKI and CKP treatment groups compared to controls on day 4 and day 7 (**A**) Jejunum day 4, (**B**) Colon day 4, (**C**) Jejunum day 7, and (**D**) Colon day 7.

GO enrichment analysis was used to compare the biological processes affected by CKI and CKP treatments and visualize the overlap of significantly enriched biological processes across the four experimental conditions using an UpSet plot. Despite the low numbers of shared DE gene between CKP and CKI in each tissue and timepoint, 13 biological processes were found to be shared across all four conditions, suggesting a set of core pathways influenced by CKI and CKP treatments (Figure 6A). To visualize enrichment patterns across the 13 shared biological processes, a heatmap was generated to show the response to treatment indicated by GO term fold enrichment using colour intensity (Figure 6B). Both the CKI and CKP groups show noteworthy enrichment in the terms “Regulation of response to stress” and “Regulation of response to stimulus”.

**Figure 6:**
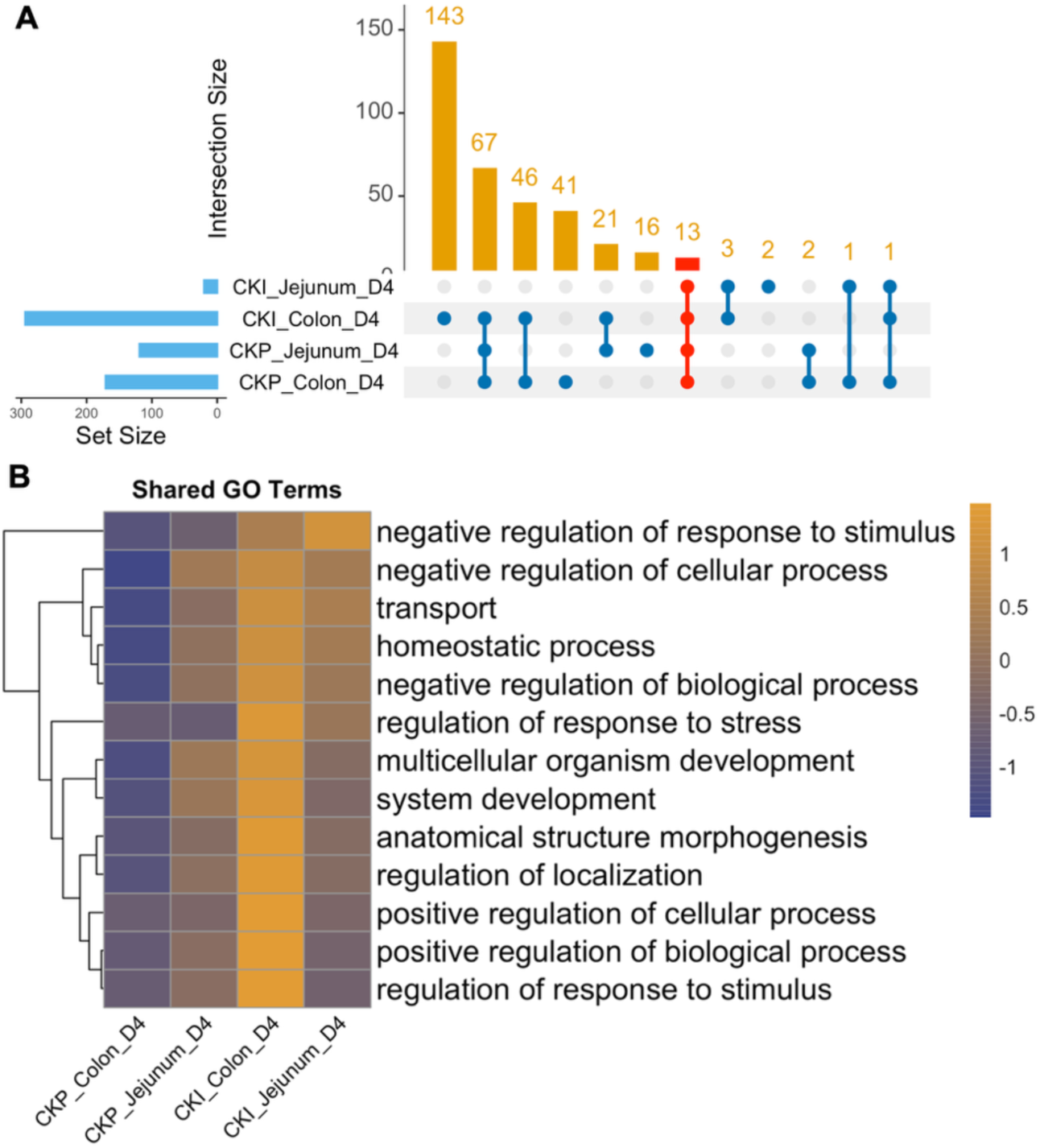
Shared and distinct GO biological process enrichment across treatment groups. (**A**) UpSet plot illustrating the intersection of enriched GO biological process terms across four comparisons. Vertical bars represent the number of overlapping GO terms, while horizontal bars indicate the total numbers of enriched GO terms in each comparison. (**B**) Heatmap displaying the fold enrichment scores of GO terms enriched in common across four treatment groups. Colours represent standardized fold enrichment values. Intensity reflects the relative DE genes fold change enrichment within a given GO term.

## 4. Discussion

This study revealed the ability of CKP and CKI to reduce diarrhoea severity and mitigate weight loss in a rat model of radiation-induced GIM, indicating their therapeutic potential. These findings, consistent with our previous study showing peak diarrhoea on day 7 post-irradiation and efficacy of CKI at 2–3 mL/kg, confirm the reproducibility of the GIM model and protective effects of CKI at 2.5 mL/kg (Harata-Lee et al., 2022). These results provide a foundation for exploring the molecular mechanisms underlying effects of these drugs through transcriptome analysis.

Transcriptome analysis revealed CKP and CKI regulate distinct and overlapping molecular pathways during the early phase of GIM. Based on the DE genes within the GIM-related panel indicates that CKP predominantly modulates inflammatory and innate immune responses in the colon, promoting early immune activation and mucosal repair. Specifically, TLR9 and TLR2 play essential roles in innate immunity, TLR9 is also known to be important in regulating pro-inflammatory cytokine production and TLR2 is critically involved in maintaining intestinal barrier integrity (Ávila et al., 2022; Cario et al., 2007). The upregulation of TNF and IL-1β along with their downstream activator of inflammation, NF-kB suggests that CKP facilitates early immune activation and mucosal repair (Sun, 2017). Additionally, the increased expression of BCL2, which inhibits apoptosis, indicates that CKP may help balance cell survival and programmed cell death, potentially minimizing excessive epithelial injury (Keefe et al., 2004). We also checked the DE gene panels generated for CKI groups, these genes enhance tissue regeneration, antioxidant defences, and anti-inflammatory signalling, particularly in the colon. The upregulation of CXCR2 and IL22 promote epithelial regeneration and wound healing, while the elevation of SOD3, an antioxidant enzyme, highlights CKI’s ability to counteract oxidative stress (Espinal et al., 2022; Sandeep and Nair, 2012; Shen et al., 2018). Additionally, the increased expression of IL4 receptor in CKI-treated samples indicates potential modulation of anti-inflammatory pathways through IL-4 signalling, which is known to reinforce epithelial barrier function and suppress excessive inflammation (Hertati et al., 2020). These findings demonstrate that during the onset of GIM, both treatments activate immune responses in the colon, but in distinct ways. CKP stimulates a strong inflammatory and regenerative response, while CKI integrates immune activation with antioxidative and barrier-enhancing effects.

Within the selected subset of DE genes associated with mucositis pathology, both CKP and CKI treatments shifted from upregulating to downregulating gene expression within their respective mucositis gene lists at the late stage of GIM. In colon tissues, CKP displayed a notable reversal in gene regulation compared to day 4. Previously upregulated genes such as *Cxcr2*, *Il22*, and *Csf2* in the colon shifted to a downregulated state. *Cxcr2* and *Il22* are known to promote epithelial regeneration during acute injury (Espinal et al., 2022; Shen et al., 2018), their suppression at this stage could lead to a reduction in regenerative demand as healing progresses. Similarly, *Csf2* (colony-stimulating factor 2) encodes a cytokine that activates eosinophils, driving production of pro-inflammatory mediators like IL-23, TNF, and IL-13, which contribute to chronic intestinal inflammation (Griseri et al., 2015). Additionally, CKP also suppressed *Itk* (IL-2 inducible T-cell kinase) in both tissues. These findings indicate that during the late stages of mucositis, CKP helps prevent excessive inflammatory responses. Another key observation was the downregulation of *Irak4* in both colon and jejunum following CKI treatment. As Irak4 is a central mediator in TLR and IL-1 receptor signalling pathways, its suppression by CKI at the later stage helps dampen excessive inflammatory responses, thereby promoting mucosal recovery and homeostasis (Garcia-Manero et al., 2024; Huang et al., 2024).

GO term analysis highlighted the enrichment of key biological processes in response to CKP and CKI treatments, particularly during the early stages of GIM. “Regulation of Response to Stress” was enriched in both CKP-treated colon and CKI-treated jejunum on day 4, with a higher number of DE genes compared to other terms, “Regulation of Response to Stimulus” was also observed in both treatments. Those processes contribute to the management of oxidative stress in intestinal tissue enhancing its ability to manage oxidative stress, which is a key driver of mucositis initiation and progression (Dong et al., 2020; Ong et al., 2010). The hallmark in early-stage mucositis is the accumulation of oxidative damage and excessive inflammatory signalling, leading to epithelial injury and ulcer formation. More genes associated with these GO terms indicate that CKI and CKP promote cellular defence mechanisms, potentially reducing inflammation-associated damage and enhancing intestinal resilience against GIM related stress factors. Both CKP treated jejunum and CKI treated colon also exhibited significant DE genes in “Response to Wounding” on day 4, refers to processes like inflammation control and tissue recovery that are essential for mucosal ulcers healing. Damage to DNA caused by radiation can be viewed as a form of wounding that is an initial pathological step of GIM, responses to such damage involve inflammatory activation, extracellular matrix remodelling, and epithelial regeneration (Sonis, 2004a). Genes uniquely regulated by CKI in the jejunum were enriched in pathways including “Response to Radiation” and “Positive Regulation of Response to DNA Damage Stimulus” highlights its potential to mitigate cellular damage and support DNA repair (Goldstein and Kastan, 2015; Treister and Sonis, 2007; Zhao and Robbins, 2009). Additionally, the significant enrichment of the “Cell Migration” term in CKI treated colon (day 4) highlights its potential involvement in epithelial regeneration by stimulating cellular movement and extracellular matrix remodelling. CKI may improve tissue remodelling and radiation resistance in a specific way.

KEGG pathway analysis revealed that CKP primarily modulates inflammatory signalling through the NF-κB and TNF pathways to attenuate mucositis-related inflammation and promotes mucosal repair while CKI acts by targeting pathways critical for cell survival and immune regulation, as evidenced by the enrichment of the PI3K-Akt and JAK-STAT signalling pathways. Under the CKP treatment, NF-κB signalling pathway was significantly enriched in the colon. As a main regulator of over 200 genes involved in inflammation, apoptosis, and cell survival, NF-κB regulates mucosal responses to injury. The activation of downstream pathways such as MAPK and COX-2 further highlights its interference in mucositis progression (Sonis, 2004b). This pathway has antiapoptotic functions in radiation induced tissue responses (Sonis, 2002). Additionally, the observed enrichment of the TNF signalling pathway points to effect of CKP on inflammatory cascades and immune modulation. Among the DE genes which were significantly downregulated in the CKP treatment group compared to the control group (Figure S1A), SOCS3 emerged as a key regulator. As a negative feedback inhibitor of cytokine signalling, SOCS3 is overexpressed in inflamed mucosa and has been implicated in tissue repair by limiting excessive inflammatory responses (Rigby et al., 2007; Thagia et al., 2015; White et al., 2011). Similarly, TLR4 is a critical mediator of mucosal immune responses which plays a pivotal role in initiating the cytokine cascade during mucositis onset. The TLR4/NF-κB axis is particularly relevant in radiation-induced mucosal damage, suggesting that the modulatory effects of CKP on these pathways may contribute to its therapeutic potential in mucositis management (Zhou et al., 2024).

CKI treatment led to significant DE gene enrichment of the PI3K-Akt signalling pathway in colon tissue. This pathway plays a crucial role in regulating cell survival and metabolism, particularly through mediating the production of inflammatory cytokines such as TNF-α and IL-1β (Yi et al., 2017). This observation indicates CKI’s potential for mitigating mucosal injury and supporting epithelial homeostasis. One of the DE genes, *Osm*, found involved in this pathway exhibited inhibited in colon (Figure S3B). As an inflammatory amplifier, OSM enhances disease chronicity by promoting stromal cell-mediated secretion of cytokines, chemokines, and adhesion molecules. Notably, its receptor OSMR is predominantly expressed on stromal cells in ulcerative colitis, driving immune cell infiltration and sustained inflammation (Hermanns, 2015; West et al., 2017). In this pathway, CKI treatment was also associated with reduced expression of gene *Epha2* which is a critical mediator of radiation-induced vascular injury (Figure S1B). Endothelial and epithelial cells exhibit increased EPHA2 phosphorylation upon radiation exposure, exacerbating inflammation and mucosal damage. Given that EPHA2 inhibition has been linked to attenuated vascular injury and inflammation, its downregulation following CKI treatment suggests a potential mechanism for mitigating mucositis severity and promoting mucosal integrity (Kim et al., 2020). In addition to PI3K-Akt signalling, DE genes also significantly clustered in the JAK-STAT signalling pathway in jejunal tissue under the treatment of CKI. This pathway is important in transducing proinflammatory cytokine signals and is highly upregulated in the intestinal mucosa of patients with active ulcerative colitis (Salas et al., 2020). *Stat3* was notably downregulated following CKI treatment in this pathway (Figure S1C). STAT3 is rapidly upregulated in intestinal epithelial cells during experimental colitis and mucosal injury models, where it drives the expression of cytokines such as IL-22, IL-6, and IL-11 to promote epithelial regeneration (Neufert et al., 2010). The observed suppression of STAT3 in CKI-treated tissues potentially mitigating excessive inflammatory responses while preserving regenerative capacity.

Secondary comparison of DE genes showed a low proportion of shared DE genes between CKP and CKI, however both overlapped on stress-adaptive processes (Zhao and Robbins, 2009). By examining the DE genes clustered within these two GO terms, MRN Complex Interacting Protein (MRNIP) was identified. This gene serves as a key regulator in mediating recombinational repair following DNA double-strand breaks (Wang et al., 2022).

Together, these findings suggest that CKP and CKI exert their therapeutic effects through different mechanisms: CKP is designed as an orally taken treatment, and the major compounds such as matrine, oxymatrine and Sophocarpine are absorbed by the intestine thus, functional compunds can work directly on local inflammation. This direct interaction with the gastrointestinal surface likely results in higher local concentrations of its active compounds, helping its ability to act against local inflammation. For instance, CKP mainly modulates early inflammatory responses via NF-κB and TNF signalling pathways, whereas CKI is an injectable formulation. The compounds enter the blood stream and work systemically either directly or by their metabolites. This likely enables CKI to exhibit broader effects on cellular processes, such as cell survival and regeneration through pathways like PI3K-Akt and JAK-STAT, which are critical for late-stage recovery from GIM. According to these distinct molecular mechanisms, a complementary therapeutic strategy combining CKP and CKI could be considered in future for treating GIM, optimising their respective features in targeting different stages of the disease. However, a limitation of this study is that we did not perform experimental validation of the pathways or genes identified. Future work could involve designing *in vitro* and *in vivo* experiments to validate their roles and further explore how CKP and CKI exert their therapeutic effects.

## 5. Conclusion

CKP is a new form of kushen/baituling extract; its effectiveness against mucositis was evaluated in a rat model of radiation-induced GIM, together with CKI. Both formulations have therapeutic potential most likely through regulation of inflammation, stress responses and repair processes. CKP primarily acts early by regulating innate immunity and inflammatory pathways (e.g. NF-κB) to reduce inflammation and promote mucosal repair. In contrast, CKI acts later by enhancing antioxidant defence and tissue remodelling, through pathways like PI3K-Akt and JAK-STAT, which support long-term recovery of the intestinal barrier. These differences are attributable to their compositions and administration routes, and highlight their potential as complementary treatments.

## Data availability statement

The datasets generated during the current study are available from the corresponding author on reasonable request.

## Acknowledgments

The authors grateful to the Bioresources team at the South Australian Health and Medical Research Institute for their technical guidance and animal care support.

## Credit authorship contribution statement

Yan Zhou: Writing – original draft, Formal analysis, Investigation, Software. Yuka Harata-Lee: Conceptualization, Writing – review and editing, Investigation. Zhipeng Qu: Writing – review and editing, Data curation, Investigation, Software. Hanyuan Shen: Investigation, Methodology. Yameng Zhang: Resources, Methodology. Xiumei Duan: Resources, Methodology. David L. Adelson: Conceptualization, Writing – review and editing, Supervision.

---

**Supplementary Figure 1:**
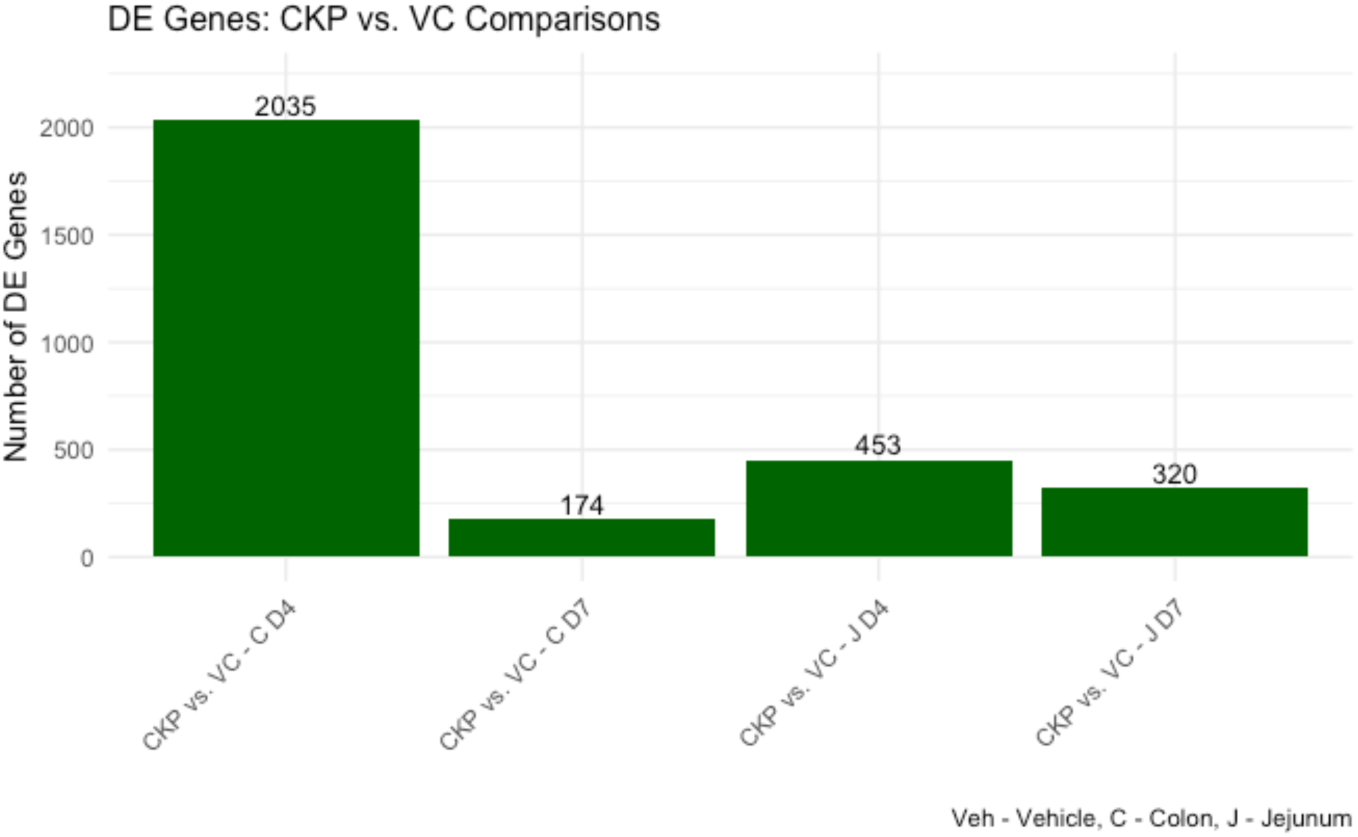
Number of DE genes in CKP comparisons. Bars represent the numbers of DE genes (P < 0.05) identified in CKP versus vehicle control comparisons at different time points and tissues. The figures above each bar indicate the numbers of DE genes in each comparison.

**Supplementary Figure 2:**
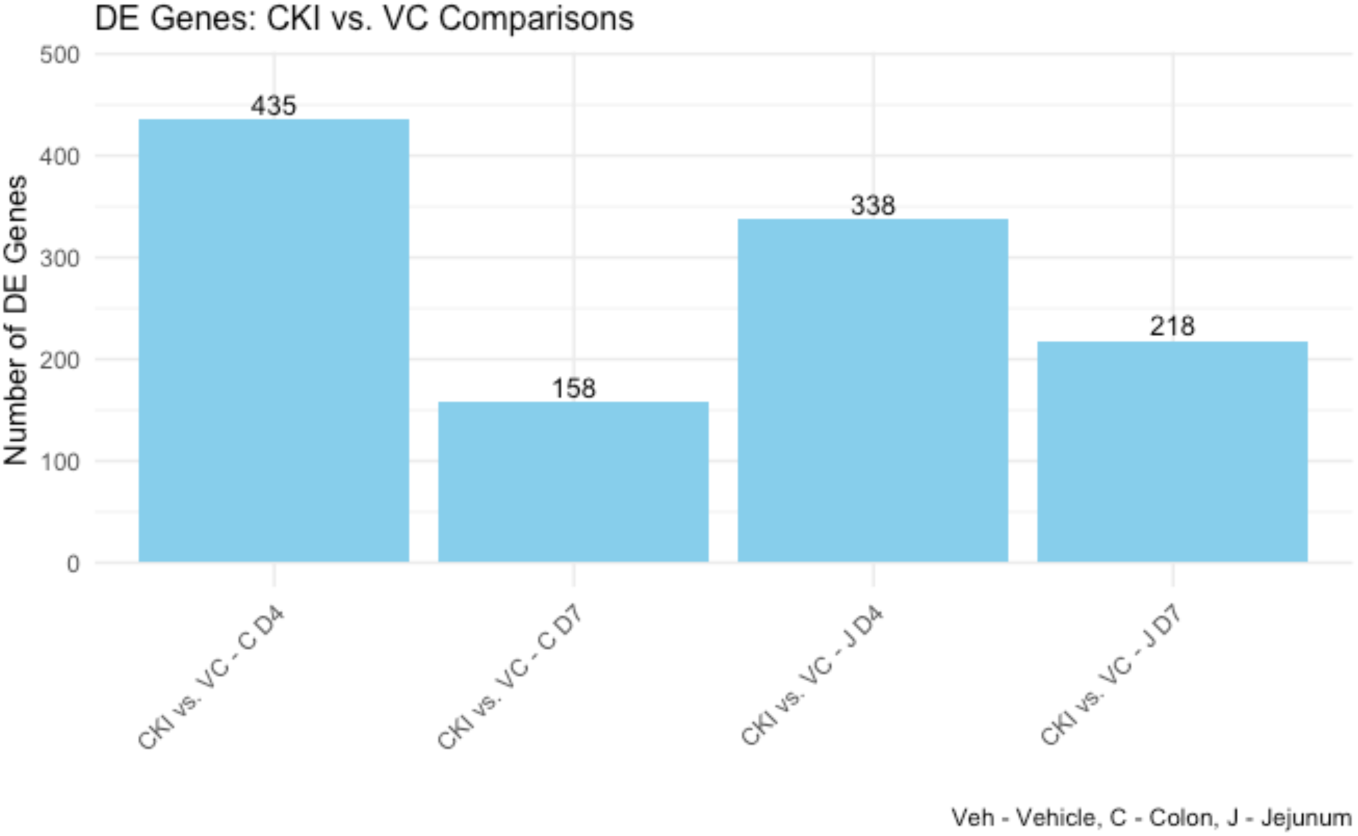
Number of DE genes in CKI comparisons. This bar plot represents the number of DE genes (P < 0.05) identified in CKI versus vehicle control comparisons at different time points and tissues. The figures above each bar indicate the numbers of DE genes in each comparison.

**Supplementary Figure 3:**
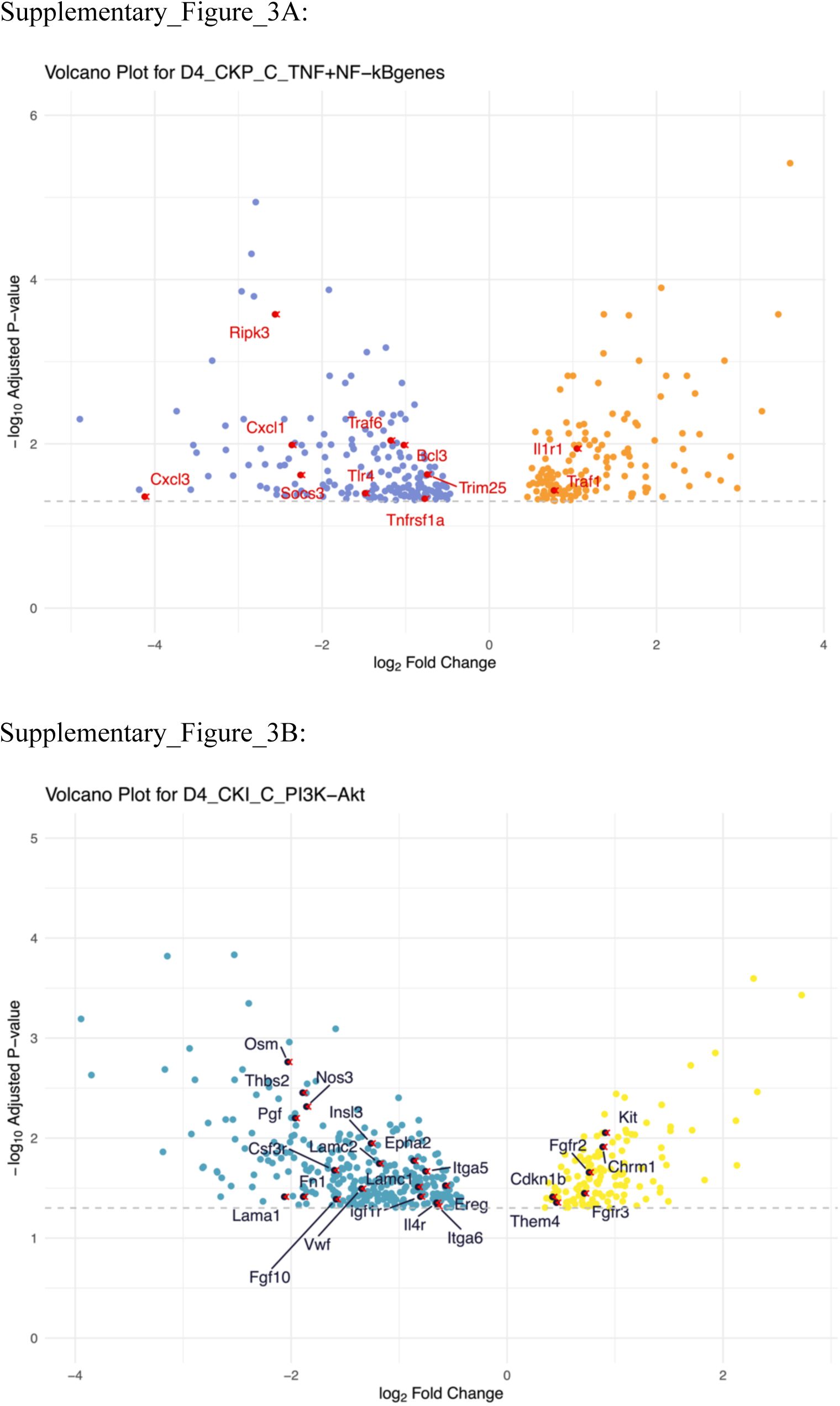
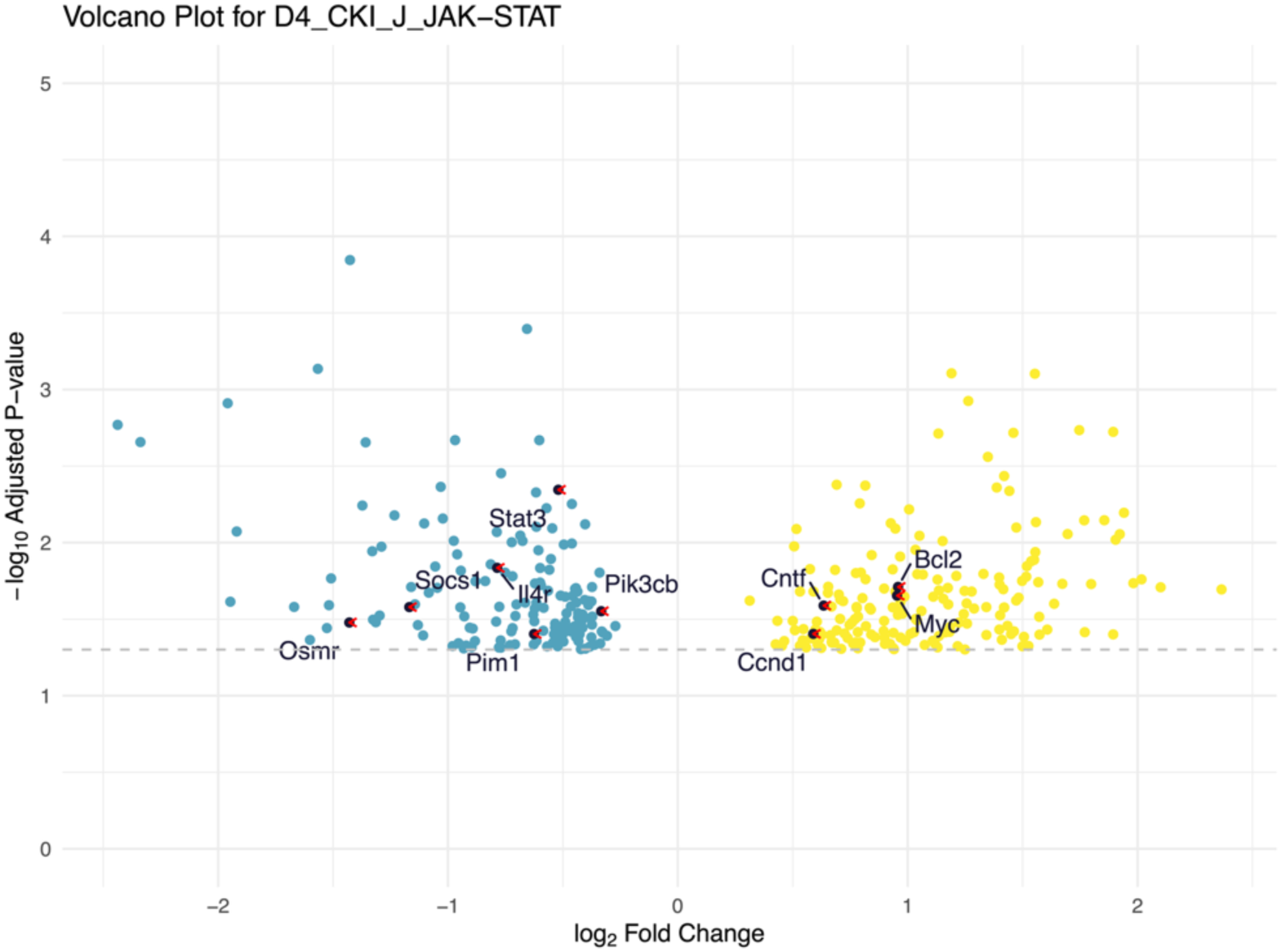
Volcano plots for DE genes in various signalling pathways on day 4 post-irradiation. (A) DE genes from the NF-κB and TNF signalling pathway in CKP treated colon tissues. (B) DE genes from the PI3K-Akt signalling pathway in CKI treated colon tissues. (C) DE genes from the JAK-STAT signalling pathway in CKI treated jejunum tissues. Those genes with adjusted p-values less than 0.05 are considered significantly differentially expressed. The x-axis represents the log2 fold change, and the y-axis represents the -log10 of the adjusted p-value. Horizontal dashed lines indicate the significance threshold (adjusted p-value < 0.05)

